# Graph-based modeling of multiparametric MRI deciphers molecular states of high-grade glioma invasion with prognostic implications

**DOI:** 10.64898/2026.06.18.733053

**Authors:** Matthew J. Flick, Matthew Kenaston, Srijon Sarkar, Gabe M. LaFond, Ian Hart, Gina Mazza, Justin Cramer, Bernard R. Bendok, Ali Turkmani, Chandan Krishna, Richard Zimmerman, Jonathon Parker, Jing Li, Kliment Donev, Krishna Bhat, Leslie C. Baxter, Yuxiang Zhou, Chad. C. Quarles, David Craig, Antonio Iavarone, Shannon Fortin Ensign, Michele Ceccarelli, Kasthuri Kannan, Nhan L. Tran, Leland S. Hu

## Abstract

The infiltrative, non-enhancing margin of IDH wildtype high grade glioma (IDHwt HGG) harbors distinct molecular programs that drive invasion and therapeutic resistance, yet remains largely unevaluable by conventional tissue sampling approaches and by conventional imaging. Here we show that this invasive architecture is encoded within multiparametric MRI (mpMRI) feature relationships and can be decoded using a graph-based framework trained on multiregional image-localized biopsies. Across 134 spatially matched biopsy–imaging pairs from 35 patients with primary IDHwt HGG (29 glioblastomas (GBM) and 6 non-glioblastoma HGGs), unsupervised graph community detection identifies two imaging-defined clusters that localize to invasive tumor regions without molecular supervision. Transcriptomic profiling associates these clusters with neuronal (NEU) and glycolytic-plurimetabolic (GPM) molecular programs.

Building on this framework, a graph convolutional network (GCN) accurately predicts NEU and GPM transcriptional states in independent training and validation cohorts and significantly outperforms conventional convolutional neural networks. Applied to whole-tumor mpMRI volumes, the trained GCN generates spatially resolved probability maps that quantify the distribution and relative burden of NEU and GPM programs across both MRI contrast-enhancing and non-enhancing invasive regions. These imaging-derived molecular maps stratify patients by overall survival. Increased GPM burden is associated with poorer survival, consistent with the aggressive behavior associated with mesenchymal-like transcriptional programs in IDHwt HGG. In contrast, increased NEU burden is associated with improved survival, identifying a previously unrecognized imaging-derived prognostic biomarker that was not detected by biopsy-based molecular classification alone. Together, these findings establish a graph-based imaging framework for spatially resolved molecular classification of invasive IDHwt HGG and demonstrate that whole-tumor molecular state architecture carries prognostic information beyond conventional tissue sampling.

## Introduction

IDH-wildtype high-grade gliomas (IDHwt HGG), most commonly glioblastoma (GBM), are aggressive primary brain tumors in adults. Molecular heterogeneity and diffuse tumor invasion are hallmark features that drive resistance, recurrence, and poor survival [1–3]. Standard surgical management is guided by conventional magnetic resonance imaging (MRI) and targets the contrast-enhancing (CE) tumor core for resection. However, this approach invariably leaves behind residual disease within the surrounding non-enhancing (NE) margins, which are subsequently treated with adjuvant chemoradiation but ultimately serve as a major source of recurrence [4, 5]. Importantly, these invasive regions are not routinely sampled during surgery and therefore remain largely unevaluated by molecular characterization [6–10]. Consequently, the biological programs and therapeutic susceptibilities underlying the tumor compartment most responsible for recurrence remain poorly defined [4, 11, 12].

Recent transcriptomic studies of IDHwt HGG, including GBM, have shown that tumors comprise distinct functional states shaped by interactions between malignant cells and the local tumor microenvironment. Pathway-based classification frameworks identify four major states distributed along metabolic and neurodevelopmental axes: glycolytic/plurimetabolic (GPM), mitochondrial (MTC), proliferative/progenitor (PPR), and neuronal (NEU) [13–15]. These states exhibit marked intratumoral heterogeneity and spatial organization. While PPR predominates within the CE tumor core, NEU and GPM states are enriched within invasive tumor margins, where they reflect distinct biological programs of invasion [4]. The NEU state is characterized by synaptic integration and quiescent tumor–neuron networks, whereas the GPM state is associated with immune and myeloid cell enrichment, mesenchymal-like programs, and an immunosuppressive microenvironment [13]. Consistent with these biological differences, mesenchymal-like GPM states are associated with poor outcome across multiple patient cohorts, whereas NEU states exhibit less aggressive phenotypes in preclinical models [15–19]. Notably, both states are more prevalent at recurrence, supporting a direct contribution of invasive tumor ecosystems to disease progression and therapeutic resistance [20–23].

The inability to routinely sample invasive tumor margins has motivated efforts to interrogate tumor biology using noninvasive imaging approaches, particularly advanced multi-parametric MRI (mpMRI) [24]. Quantitative biomarkers derived from perfusion, diffusion, and structural MRI capture complementary information related to vascularity, cytoarchitecture, tissue microstructure, cellular packing, and edema across spatially distinct tumor regions [25–38].

These imaging features have increasingly been combined with machine learning and deep learning approaches to predict molecular characteristics of HGG, particularly through image-localized biopsy studies designed to capture region-specific imaging signatures in the context of intratumoral heterogeneity [10, 39–41]. However, predictive performance remains constrained by the limited size of image-localized biopsy datasets and by analytical frameworks that treat biopsies as independent observations, rather than leveraging the biological relationships among spatially distributed tumor regions [42, 43]. In addition, many existing approaches rely on high-dimensional radiomic texture features that are difficult to interpret in the context of the underlying biology.

Graph neural networks offer a principled solution by explicitly representing samples as nodes and relationships between them as edges, enabling learning from relational structure rather than independent feature vectors [44–46]. Such representations are well matched to the ecology of IDHwt HGG invasion, where biological similarity is not constrained by anatomical proximity [12]. Graph-based approaches also accommodate sparse and irregular sampling, making them particularly suitable for image-localized multi-regional biopsy datasets, in which molecular ground truth is available only at a limited number of spatial locations within a continuously heterogeneous tumor.

Here, we develop a graph-based framework that integrates image-localized biopsies, transcriptomic profiling, and multi-parametric MRI to resolve molecular heterogeneity within invasive IDHwt HGG [47–49]. Briefly, imaging features are organized into hierarchical "imaging events" representing groups of samples with similar radiographic characteristics across multiple scales. These events form the nodes of a graph, while shared sample membership defines the edges between them, enabling the model to learn from relationships among imaging patterns rather than from independent feature vectors alone. We show that unsupervised graph-based clustering identifies biologically meaningful imaging communities enriched for NEU and GPM molecular states. Building on this structure, we develop a graph convolutional network (GCN) classifier that predicts molecular state directly from MRI and substantially outperforms a matched CNN benchmark based on biopsy-centered imaging patches, demonstrating the importance of relational imaging information beyond local spatial features [47, 48]. We further generate whole-tumor molecular state maps that quantify the distribution and burden of NEU and GPM states across whole tumor volumes, enabling patient stratification by survival. Increased GPM burden is associated with poor outcome, consistent with the aggressive behavior of mesenchymal-like programs in IDHwt HGG. Importantly, we identify a previously unrecognized association between increased NEU burden and improved overall survival, establishing the NEU invasive state as an imaging-derived prognostic biomarker not captured by conventional biopsy-based molecular subtyping. Together, these findings establish a graph-based imaging framework that spatially resolves IDHwt HGG molecular heterogeneity and enables noninvasive molecular state classification of invasive tumor ecosystems.

## Results

### Regional context fundamentally alters the biological interpretation of MRI biomarkers in IDHwt HGG

A fundamental challenge in HGG imaging biomarker research is that biological interpretation of an imaging features depends on where within the tumor it is measured. To systematically evaluate this context-dependence, we analyzed 134 spatially matched biopsy-imaging pairs from 35 primary IDHwt HGG (29 GBM, 6 other HGG) tumors profiled with 28 multi-parametric MRI (mpMRI) features and pathway-based transcriptional state subtype scores. Global associations between imaging features and molecular programs were weak and heterogeneous across the full cohort (Supplementary Figure 1A), consistent with the biological complexity of HGG and the limitation of analyses that pool anatomically distinct regions.

In contrast, region-stratified analyses revealed widespread directional reversals in imaging-biology relationships. Multiple MRI features exhibited significant opposing associations with NEU state scores between the contrast-enhancing (CE) core and non-enhancing (NE) infiltrative margin (Figure 1A-B). For example, normalized K2 (nK2) acquired with consensus Dynamic Susceptibility Contrast (DSC) MRI (30° flip angle, FA30) demonstrated a strong inverse relationship with NEU scores in NE samples but an attenuated and directionally opposite relationship in CE samples [4]. This phenomenon was not limited to nK2 but extended across a broad range of mpMRI features, indicating that the biological meaning of imaging biomarkers is strongly dependent on regional context.

**Figure 1.**
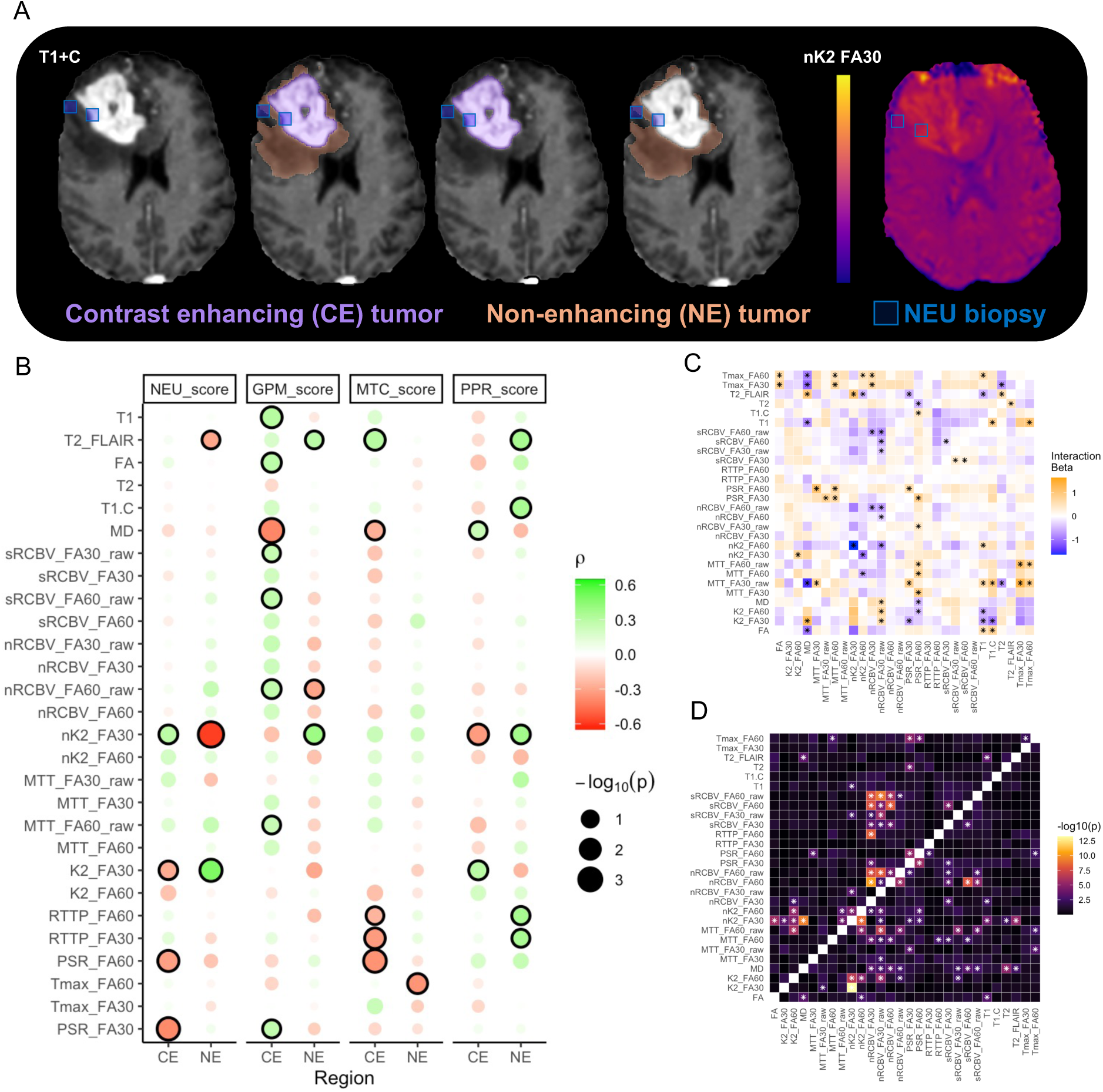
Tumor region and biological subtype influence mpMRI features. **A.** Representative contrast-enhancing (CE) and non-enhancing (NE) tumor regions showing exemplary overlay of tumor biopsies of Neuronal subtype collected from NE and CE-NE boundary, with regionally variable nK2 signal. **B.** Feature-subtype score Spearman correlations grouped by tumor region (NE/CE). Significant feature-subtype score relations (p<0.05) denoted with a black outline. **C.** Interaction of tumor region (NE/CE) with multiparametric feature-feature associations in linear mixed effect models. **D.** Significance of tumor subtype by ANOVA with respect to multiparametric feature-feature associations accounting for nested patient effects. *Significant correlations or model effect terms (FDR<0.05).

Intraclass correlation analysis revealed substantial between- patient variability for multiple imaging metrics (Supplementary Figure 1B), supporting the use of mixed-effects modeling to accounts for repeated sampling within patients. All subsequent analyses therefore incorporated patient identity as a random effect (Figure 1C-D, Supplementary Figure 1C). Beyond individual feature associations, pairwise inter-feature imaging relationships varied markedly as a function of both anatomical region and molecular state. Metrics including nRCBV, nK2, and T1+C exhibited pronounced regional sensitivity, while sRCBV and K2, among others, remained comparatively stable across regions (Figure 1C). Molecular state exerted an additional modulatory effect on feature–feature relationships, with nRCBV and PSR showing the strongest state-dependent variability (Figure 1D, Supplementary Figure 1D-E).

Together, these findings demonstrate that the mpMRI feature landscape is jointly shaped by anatomic region, molecular state, and their interactions. The resulting context dependence cannot be adequately represented by conventional feature-centric approaches and motivates a relational modeling framework that explicitly captures biological relationships among spatially distributed tumor regions.

### Unsupervised graph community detection recovers molecular programs of HGG invasion from imaging alone

Having established that inter-regional imaging relationships encode biologically meaningful information, we next asked whether organizing mpMRI data as a relational graph could recover the invasive biology of IDHwt HGG without molecular supervision. We constructed a hierarchical imaging event graph from 28 mpMRI features across 134 image-localized biopsies by representing multiscale imaging clustering events, derived from dendrogram partitions generated through per-feature hierarchical clustering, as graph nodes and shared sample membership between events as edges (Figure 2A). Louvain modularity optimization across 1,000 independent iterations identified 1,042 stable graph communities, many of which were enriched for invasive NE samples.

**Figure 2.**
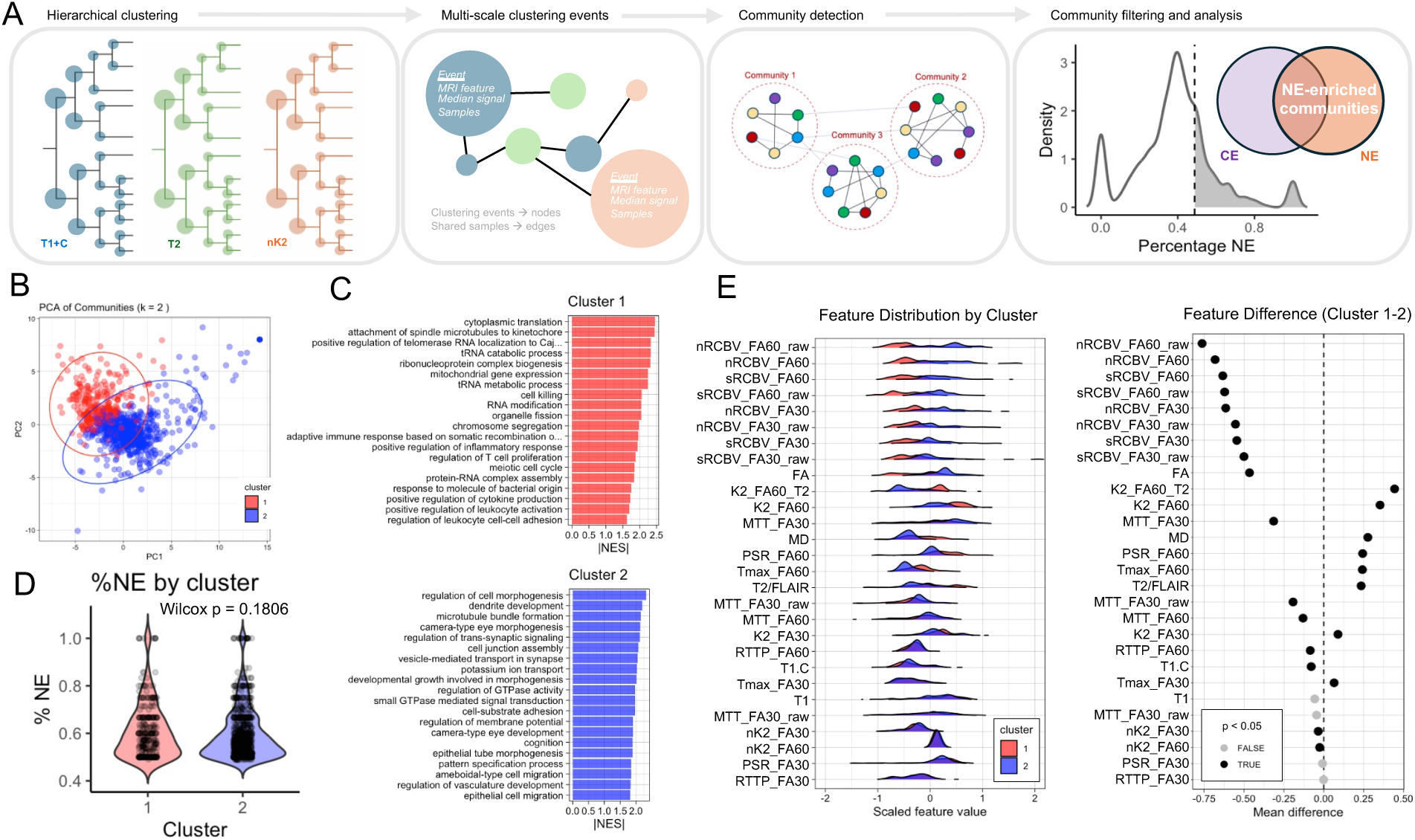
Graph-based mpMRI communities reveal tumor margin-enriched neuronal and immune states. **A.** Representative pipeline for initial analysis and selection of NE-enriched multi-resolution mpMRI communities generated through a hierarchically clustered, graph network-based approach. **B.** PCA of selected communities using median imaging signal aggregated at the community level with annotation of two dominant community clusters. **C.** Gene-set enrichment analysis for differentially expressed genes generated using frequency-weighted distributions of samples in cluster 1 versus cluster 1. **D.** Wilcox test for difference of %NE composition in communities in cluster 1 versus cluster 2. **E**. Relative imaging feature distributions for cluster 1 and cluster 2 aggregated using median signal per feature at the community level (left) and difference between mean imaging feature signal for cluster 1 versus cluster 2, reflecting feature signal differences at the cluster level that incorporate a heterogeneous mixtures of NE and CE samples at the community level.

Aggregation of community-level imaging signatures resolved two dominant imaging-defined clusters (Figure 2B). Transcriptomic pathway enrichment analysis revealed a striking biological separation between these clusters: one was enriched for immune signaling, glycolysis, hypoxia, and metabolic reprogramming, consistent with the GPM state, whereas the other was enriched for neuronal differentiation, synaptic organization, neurotransmitter signaling, and morphogenic programs characteristic of the NEU state (Figure 2C). Notably, this separation emerged entirely without molecular supervision, indicating that graph topology derived from imaging relationships alone is sufficient to recover biologically meaningful invasive phenotypes.

Importantly, the two imaging-defined clusters did not differ significantly in their CE versus NE sample composition (Figure 2D), demonstrating that graph-derived community identity is not simply a surrogate for anatomical location. Both GPM-associated and NEU-associated communities contained mixed CE and NE samples, indicating that community phenotype is orthogonal to the CE/NE anatomical boundary that underlies conventional radiologic and surgical classification. Among discriminating features, perfusion-derived rCBV metrics showed the strongest separation between clusters (Figure 2E). The immune-enriched GPM state exhibited reduced perfusion signatures, consistent with inflammatory remodeling of the microvasculature and reduced functional microvascular volume, a phenomenon associated with inflammation-induced vascular thrombosis and collapse [49, 50]. Together, these observations identify vascular attenuation as a distinguishing imaging feature of the GPM invasive ecosystem.

### GCNs outperform conventional deep learning for decoding invasive molecular states

Building on the unsupervised community structure, we trained supervised graph convolutional network (GCN) models to predict NEU and GPM transcriptional states directly from mpMRI features and benchmarked their performance against matched patch-based convolutional neural networks (CNNs) trained on spatially corresponding imaging patches from the same patients (Figure 3A-B; Supplementary Figure 2A-B) [13,14,20,51]. Model training used a pre-specified tumor-level cohort split, ensuring that all biopsies from a given tumor were assigned exclusively to either the training or testing set and preventing data leakage.

**Figure 3.**
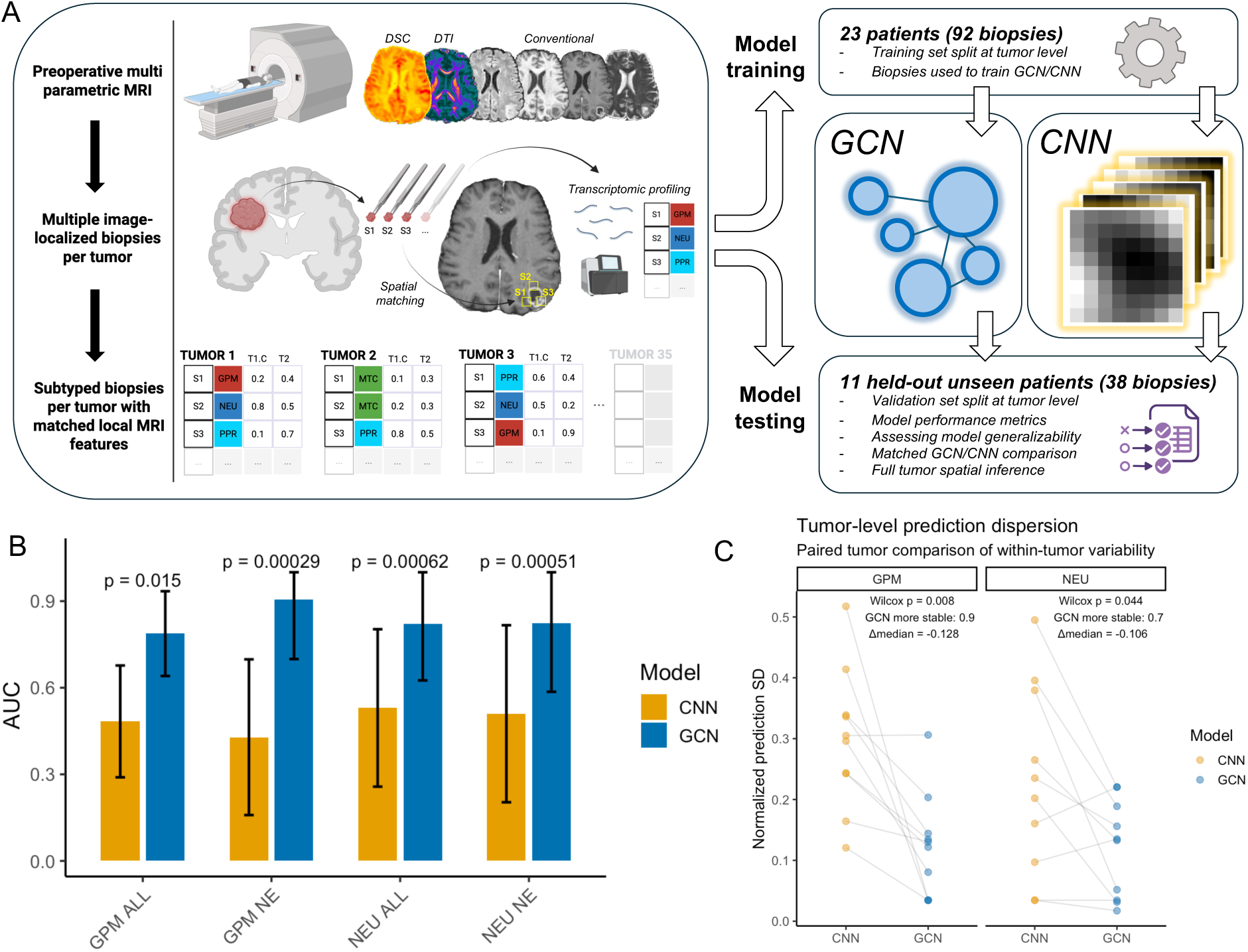
GCN outperforms CNN benchmark for biopsy level subtype classification. **A.** Pipeline for collection of image localized biopsies with matched mpMRI features with downstream patient-level training/test split for GCN and CNN model development and performance evaluation. **B.** DeLong test for GCN versus CNN AUCs on all matched held out test biopsies (n = 38 biopsies, 11 patient tumors) and of NE samples (n = 23 biopsies) only with 95% confidence intervals. DeLong p-values represent AUC difference using covariance-aware paired statistics on the matched samples. **C.** Tumor-level prediction dispersion, defined as the within-tumor standard deviation of normalized CNN and GCN scores, shown as paired tumor-wise comparisons across NEU and GPM cohorts. Paired lines connect model-specific dispersion within each tumor.

GCNs substantially outperformed CNNs across both classification tasks. For NEU prediction, the GCN achieved an AUC of 0.82 compared with 0.52 for the CNN (paired DeLong test, p = 0.00062). For GPM prediction, the GCN achieved an AUC of 0.79 compared with 0.49 for the CNN (p = 0.015) (Figure 3B). Performance remained high within non-enhancing (NE) invasive samples, where the GCN achieved AUCs of 0.82 for NEU and 0.90 for GPM, significantly exceeding CNN performance (0.50 and 0.42, respectively; p < 0.001). These findings indicate that graph-based models retain predictive accuracy within the infiltrative tumor compartment that is most relevant to recurrence.

GCNs also exhibited significantly lower within-tumor prediction dispersion than CNNs for both NEU (p = 0.044) and GPM (p = 0.008) classification tasks (Figure 3C), indicating greater consistency among biopsies derived from the same tumor. This observation suggests that graph-based models capture patient-level biological structure rather than local imaging variation alone. Consistent with this interpretation, mRMR-based feature selection had minimal impact on GCN performance but substantially affected CNN performance (Supplementary Figure 2C), suggesting that graph message passing and neighborhood aggregation reduce dependence on predefined dimensionality reduction.

Despite the relatively small size of the held-out test cohort (38 biopsies from 11 patient tumors), these results demonstrate that graph-based learning provides a substantial advantage for decoding molecular states from mpMRI. By leveraging relational structure among spatially distributed tumor regions, GCNs more effectively capture imaging–molecular associations underlying NEU and GPM invasion phenotypes than conventional CNN architectures.

### Whole-tumor GCN maps capture the spatial extent of invasive molecular state subtypes

To extend biopsy-level predictions to the tumor scale, we applied trained GCN models to whole-tumor mpMRI volumes and generated voxel-wise probability maps of NEU and GPM molecular states (Supplementary Fig. 3A). In brief, overlapping tumor patches were converted into imaging-event graphs analogous to those used during model training, and GCN-derived probabilities were assigned back to individual patches before being interpolated into continuous spatial maps across the tumor volume. This enabled evaluation of whether molecular state representations learned from sparse image-localized biopsies generalize across the entire tumor landscape. Predicted molecular state probabilities were extracted at known biopsy coordinates and compared with matched RNA-derived molecular state scores. Following within-tumor normalization, GCN-derived maps showed significant correlations with transcriptomic measurements for both NEU (ρ = 0.51, p = 0.003) and GPM (ρ = 0.59, p = 0.0004) (Figure 4A; Supplementary Figure 3B). Mixed-effects modeling confirmed that these associations remained significant after accounting for tumor-level random effects (NEU: β = 0.095, p = 0.021; GPM: β = 0.104, p = 0.0029), indicating that imaging–molecular concordance reflects within-tumor spatial variation rather than inter-patient differences. Consistent with these findings, RNA-defined NEU biopsies exhibited significantly higher GCN-derived NEU scores (β = 0.89 ± 0.37 SE, p = 0.023), whereas RNA-defined GPM biopsies exhibited significantly higher GCN-derived GPM scores (β = 0.52 ± 0.25 SE, p = 0.046).

**Figure 4.**
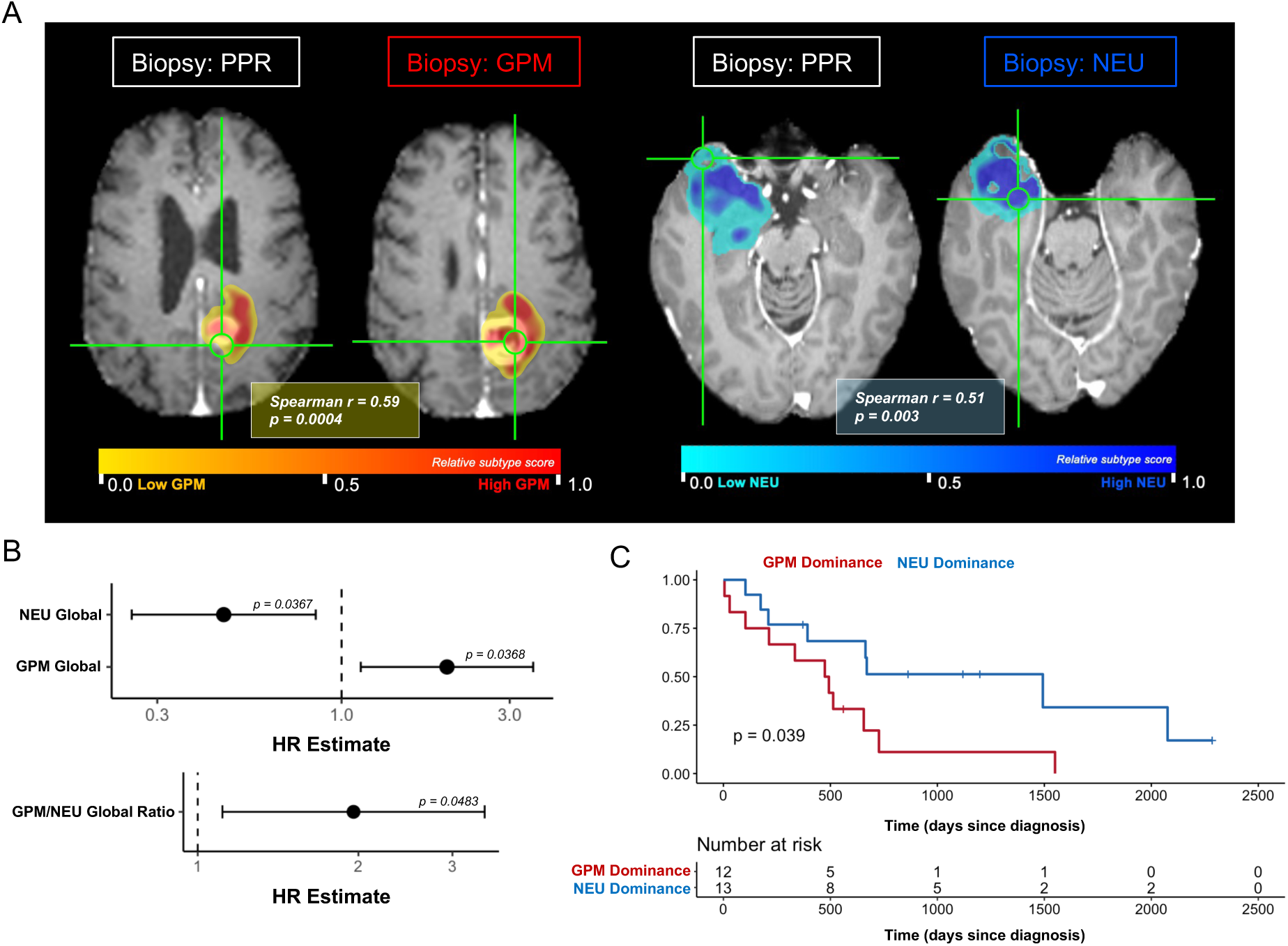
Tumor-level GCN inference recapitulates biological ground truth with survivorship delineation. **A.** Representative GCN-predicted subtype maps for two exemplary tumors from the held-out test set with annotation of confirmed biological subtype. GCN-predicted maps are of within-tumor scaled relative subtype probability scores. Spearman correlations of RNA-derived biological subtype score with values extracted from full tumor GCN-inferred subtype map at biopsied locations in held out test patient tumors (n = 8 patient tumors, 32 biopsies). **B.** Cox proportional hazards model assessment of associations between GCN-derived combined NEU enrichment (NEU Global), combined GPM enrichment (GPM Global), and the global GPM/NEU dominance ratio with overall survival, adjusting for age and non-enhancing tumor volume fraction (n = 25 patients). **C.** Kaplan–Meier analysis stratified tumors by median global GPM/NEU dominance (n = 25 patients).

Because NEU and GPM represent opposing invasion programs, we next examined a continuous molecular state balance axis defined as the difference between NEU and GPM scores. This measure demonstrated substantially stronger imaging–molecular agreement than either state alone (ρ = 0.71, P = 8.4 × 10⁻⁶; Supplementary Figure 3B). Mixed-effects modeling confirmed a significant association between MRI-derived and RNA-derived state balance (β = 0.158, P = 4.4 × 10⁻⁶). A significant state balance × region interaction (β = −0.115, P = 0.017) further demonstrated that imaging–molecular relationships differ between CE and NE compartments, recapitulating the regional context dependence identified in Fig. 1.

At the individual patient level, median within-tumor correlations between GCN predictions and transcriptomic measurements were moderate for both NEU (r = 0.54) and GPM (r = 0.58), with substantial variation across tumors (Supplementary Figure 3C). Together, these findings demonstrate that GCN-derived molecular state maps generalize beyond sampled biopsy locations and capture biologically meaningful spatial variation across the entire tumor.

### Whole-tumor GCN maps of molecular state independently predict overall survival

We next asked whether GCN-derived whole-tumor molecular state maps carry prognostic information beyond conventional clinical variables [15]. In a cohort of 25 patients with the complete panel of mpMRI imaging and outcome data (23 GBM, 2 IDHwt HGG), multivariable Cox proportional hazards models adjusting for age and non-enhancing (NE) tumor volume demonstrated that increased GPM burden independently predicted worse overall survival (hazard ratio [HR] 1.90, 95% CI 1.04–3.49, p = 0.037), whereas increased NEU burden independently predicted improved survival (HR 0.52, 95% CI 0.28–0.96, p = 0.037) (Figure 4B-C). A continuous molecular state dominance score, defined as the relative balance between GPM and NEU burden, was similarly associated with worse survival after covariate adjustment (HR per SD 1.85, 95% CI 1.00–3.41, p = 0.048). Consistent with these findings, Kaplan–Meier analysis dichotomized at the median dominance score demonstrated significantly worse overall survival in GPM-dominant tumors (log-rank p = 0.039; Figure 4C). Median overall survival for patients with GPM-dominant tumors was 15.9 months (483.5 days) versus 22.0 months (670 days) for patients with NEU-dominant tumors. Survival associations remained stable across multiple model specifications, including GBM-restricted analysis, and covariate adjustments (Supplementary Figure 3D). To determine whether these prognostic associations reflected whole-tumor ecosystem architecture rather than the molecular composition of sampled biopsies, we repeated the analysis using biopsy-derived molecular class fractions. In contrast to GCN-derived tumor maps, neither GPM biopsy fraction (HR 0.86, p = 0.53) nor NEU biopsy fraction (HR 1.48, p = 0.096) was independently associated with survival. A biopsy-level competition score, defined as the log-ratio of GPM to NEU biopsy fractions, was similarly non-significant (p = 0.47). These findings indicate that prognostic information is encoded in the spatial distribution and burden of molecular states across the tumor rather than in the composition of a limited set of sampled biopsies.

## Discussion

A central finding of this study is that relational patterns among mpMRI features encode biologically meaningful information about the molecular states of HGG invasion that is not readily captured by conventional imaging analyses. By representing image-localized biopsies as a graph and learning from relationships among spatially distributed tumor regions, we recover and spatially resolve the NEU and GPM pathway-based molecular state subtypes known to be enriched in invasive tumor regions. This enables generation of whole-tumor maps of molecular state burden and reveals associations between molecular architecture and patient survival.

Together, these findings suggest that clinically relevant information is encoded in the spatial distribution and burden of molecular states across the tumor and can be recovered noninvasively through imaging.

The improved performance of GCNs relative to conventional CNNs likely reflects fundamental differences in how these frameworks represent tumor heterogeneity and manage modestly sized image-localized biopsy datasets comprised of sparsely sampled tumors [51, 52]. IDHwt HGG invasion is characterized by spatially distributed molecular states that do not necessarily follow anatomical boundaries, with biologically related regions often separated in space and molecularly distinct states frequently coexisting within close proximity. Whereas CNNs learn primarily from local image neighborhoods, GCNs explicitly model relationships among biologically similar regions regardless of physical location. Consistent with this interpretation, GCN performance remained robust despite feature selection, suggesting that graph-based representations can leverage distributed biological information that is difficult to capture using feature-centric approaches alone. More broadly, these findings indicate that graph-based learning may be particularly advantageous for image-localized biopsy studies, where sparse sampling and limited cohort sizes constrain conventional machine learning and deep learning approaches [4, 10, 39–41].

An important observation is that whole-tumor imaging-derived molecular state burden demonstrated strong prognostic associations, whereas biopsy-derived subtype composition did not. While molecular profiling provides high-resolution characterization of individual sampling locations, it captures only a small fraction of the tumor ecosystem and may therefore miss biologically important patterns of spatial organization. Previous transcriptomic studies have linked mesenchymal-like GPM programs to poor outcome and aggressive disease behavior, findings that are recapitulated by our imaging-derived molecular maps [15, 18, 19, 53]. In addition, we identify a previously unrecognized association between increased NEU burden and improved overall survival. The ability to detect prognostic effects using whole-tumor molecular state maps suggests that the spatial distribution and burden of molecular states across the tumor may be more informative than localized molecular measurements alone. These findings further identify NEU burden as an imaging-derived prognostic biomarker that is not captured by conventional biopsy-based molecular classification and support the concept that ecosystem-scale organization contributes independently to clinical outcome.

The ability of graph-derived imaging communities to recover NEU and GPM phenotypes without molecular supervision further supports the concept that advanced imaging features capture emergent biophysical properties of invasive tumor ecosystems [33]. The strong separation of GPM and NEU communities by DSC-MRI perfusion biomarkers suggests that differences in microvascular organization and microenvironmental composition are major determinants of their imaging phenotype. In particular, the reduced perfusion signatures observed in immune-enriched GPM ecosystems are consistent with prior evidence that inflammatory remodeling can attenuate functional microvascular volume through vascular dysfunction and thrombosis [49, 50]. Similarly, the ability of DSC-derived nK2, a biomarker associated with cytoarchitectural heterogeneity, to distinguish molecular states in this larger cohort reinforces prior observations that advanced MRI captures biologically meaningful differences in tissue organization within invasive tumor regions [4, 38]. Together, these findings suggest that multiparametric imaging provides a noninvasive window into the biophysical organization of HGG invasion and the ecosystem-level processes that shape molecular state identity.

Several limitations should be considered. First, although this study represents one of the largest image-localized, multiregional IDHwt HGG biopsy cohorts currently available, the sample size remains modest and independent validation in larger multi-institutional cohorts will be required to establish generalizability. Second, the survival analyses should be considered hypothesis-generating and warrant prospective validation, particularly the association between increased NEU burden and improved outcome. Third, while transcriptomic profiling provides molecular ground truth for NEU and GPM states, the extent to which imaging-derived communities reflect specific cellular ecosystems and spatial architectures remains to be determined through integration with single-cell and spatially resolved molecular datasets. Finally, although GCNs substantially outperformed CNNs in this setting, future studies should evaluate graph-based approaches relative to a broader range of emerging architectures.

More broadly, these findings suggest that relational structure within imaging data provides a powerful framework for studying spatially organized biological heterogeneity in cancer. By enabling noninvasive mapping of molecular state architecture across the entire tumor landscape, graph-based imaging approaches may complement tissue profiling and provide new opportunities to investigate regional ecosystem dynamics, therapeutic response, and disease evolution in HGG and other heterogeneous solid tumors [54].

## Methods

### Patient Cohort, Data Acquisition and Transcriptome Analysis

This study was conducted under IRB approval from Barrow Neurological Institute and Mayo Clinic Arizona, with the Mayo Clinic IRB serving as the coordinating protocol; written informed consent was obtained from all participants. Analyses were restricted to treatment-naive primary IDH-wildtype HGG (29 GBM, 6 other HGG), yielding 134 image-localized biopsies from 35 primary tumors after quality-control exclusions. Of the 35 tumors included in this study, 22 (82 biopsies) were derived from previously reported cohorts [4], whereas 13 tumors (52 biopsies) are reported here for the first time. Preoperative multiparametric MRI (mpMRI), including anatomical, diffusion, and perfusion imaging, was acquired on 3-T clinical scanners and processed using previously described image registration and quantitative map generation pipelines [4]. Spatially localized stereotactic biopsies were coregistered to preoperative MRI, and biopsy-matched imaging features were extracted from standardized regions of interest as previously described [4]. RNA was extracted from frozen surgical biopsies and sequenced on the Illumina platform; reads were processed and normalized as detailed in Supplementary Methods. Transcriptional subtypes (neuronal [NEU], glycolytic/plurimetabolic [GPM], mitochondrial [MTC], and proliferative/progenitor [PPR]) were assigned using the pathway-based classifier of Garofano et al. [13], with samples below enrichment thresholds classified as non-classified (nc).

### Imaging feature–biology association analysis

To characterize relationships between mpMRI-derived imaging features and tumor transcriptional biology, we evaluated imaging–molecular associations across complementary analytical frameworks. Global associations between imaging features and transcriptional subtype scores (NEU, GPM, MTC, PPR) were assessed across all samples using two-sided Spearman rank correlations, with Benjamini–Hochberg FDR correction applied across all feature–score comparisons; correlation matrices were visualized as heatmaps, with FDR-significant associations (q<0.05) annotated. These associations were then decomposed by anatomical tumor compartment — contrast-enhancing (CE) versus non-enhancing (NE) — using region-stratified Spearman correlations and linear interaction models on z-score-standardized imaging features, to identify region-dependent shifts and directional sign-flips in feature–score relationships. To account for repeated regional sampling within patients, linear mixed-effects models with patient-specific random intercepts were used to derive region-specific (CE vs. NE) effect sizes, and intraclass correlation coefficients were computed to quantify the proportion of variance in each imaging feature attributable to inter-patient variability. Finally, pairwise mixed-effects interaction models tested whether relationships between imaging features were themselves region-dependent or subtype-dependent, with all pairwise comparisons FDR-corrected (q<0.05). For specific details, refer to Supplementary Methods

### Community Detection

Imaging features from discovery and validation cohorts were combined into a unified feature space and preprocessed by removing near-zero-variance and highly collinear features, followed by z-score normalization (details in Supplementary Methods). For each imaging feature independently, hierarchical agglomerative clustering (Euclidean distance, median linkage) was performed across all samples, and all internal dendrogram partition events were extracted to generate a multiscale representation of imaging similarity; each event was annotated with its constituent samples, median signal intensity, feature identity, and regional (CE/NE) composition. These events were imported into a Neo4j graph database as nodes, with edges connecting events that shared constituent samples, weighted by the number of shared samples. Stable higher-order imaging communities were identified using Louvain modularity optimization applied to iteratively sampled subgraphs (1,000 iterations), with results merged into a consensus community table. Consensus communities were annotated for NE/CE regional composition and characterized by median imaging-feature signatures across constituent samples, excluding small or low-diversity communities. Communities in the upper quartile of NE enrichment (>75th percentile) were defined as highly invasive, and communities with identical sample sets were collapsed. Community-level imaging signatures underwent principal component analysis (retaining components explaining ≥85% of variance) followed by Ward’s hierarchical clustering (ward.D2) in PCA space; a two-cluster solution was selected based on silhouette analysis and biological interpretability, defining two dominant axes of invasive imaging organization.

### Imaging Community–Transcriptomic Integration

The two imaging-derived community clusters identified in Section 3 were used as the basis for transcriptomic integration, linking spatial imaging organization to molecular subtype programs at the biopsy level. Each biopsy was assigned continuous cluster membership ratios based on the proportion of its graph communities belonging to each imaging cluster. Gene expression counts were normalized using TMM normalization and voom transformation (limma), and differential expression was modeled with a linear design incorporating continuous cluster ratio and sequencing batch as covariates. Genes were ranked by moderated t-statistics from the cluster ratio coefficient and analyzed via GO Biological Process gene set enrichment (gseGO, clusterProfiler); redundant pathways were collapsed using semantic similarity (rrvgo), and top enriched pathways were prioritized by adjusted p-value and normalized enrichment score to identify molecular programs associated with each imaging-defined invasive cluster. Differences in imaging-feature profiles and NE enrichment between the two clusters were assessed using two-sided Wilcoxon rank-sum tests with FDR correction, and visualized via mean scaled feature plots, ridge density plots, and violin/box plots.

### Graph Convolutional Network Modeling

Building on the imaging community framework (Section 3), we developed graph convolutional network (GCN) models to predict molecular subtype status directly from imaging-derived features, constructing separate binary classifiers for neuronal (NEU vs. non-NEU) and glycolytic/plurimetabolic (GPM vs. non-GPM) subtypes. Training and test cohorts were split at the tumor level to prevent data leakage (23 training patient tumors with 92 biopsies, 11 held out test patient tumors with 38 biopsies), and imaging features were preprocessed using the same near-zero-variance, correlation, and z-score normalization pipeline as Section 3.1, with imputation and scaling parameters derived exclusively from the training cohort (28/28 features retained). For each task, minimum redundancy–maximum relevance (mRMR) feature selection identified the top-ranked features for downstream modeling (full feature lists in Supplementary Methods); CNN and GCN models using these selected subsets were trained identically to the full-feature models to isolate the effect of feature dimensionality. Hierarchical imaging events and graphs were constructed for the NEU and GPM tasks following the pipeline in Sections 3.2–3.3, with nodes additionally annotated by subtype composition; binary node labels were assigned using a ≥90% subtype-purity threshold. Graphs were converted to PyTorch Geometric objects, with node features comprising the median imaging signal and a learned embedding of imaging feature identity, and edge weights derived from log-transformed, normalized shared-sample counts. GCN models consisted of a feature-identity embedding layer, two graph convolution layers (GCNConv) with ReLU activation and dropout, and a final binary classification layer. Training used weighted cross-entropy loss to address class imbalance, combined with a sample-consistency regularization term penalizing prediction variance across nodes derived from the same biopsy. Hyperparameters were optimized using Optuna with five-fold stratified cross-validation on training/validation nodes, selecting based on mean validation F1 score. Final models were retrained on combined training and validation nodes with early stopping, and node-level probabilities were aggregated to sample-level predictions via mean-logit aggregation followed by sigmoid transformation. Performance was evaluated using ROC-AUC, F1, and threshold-based accuracy at both node and sample levels. Independent test graphs were generated from held-out test tumors using an identical pipeline, and trained model checkpoints were applied without retraining, with performance again assessed via ROC-AUC at node and sample levels.

As a benchmark, multimodal CNN classifiers (NEU vs. non-NEU and GPM vs. non-GPM) were trained on biopsy-local multimodal MRI patches using the same patient-level train/test split as the GCN models, enabling matched comparison. CNN and GCN ROC curves on matched held-out samples were compared using paired DeLong tests. Patient tumors from the test set were included in ROC-AUC analysis if preoperative imaging was within 1-2 weeks of surgery (11 patient tumors, 38 biopsies). 1 patient tumor (4 biopsies) were excluded because imaging was performed 4 weeks before surgery, resulting in misregistration errors.

### Full Tumor ROI Analysis

To extend biopsy-level GCN predictions to whole-tumor spatial maps, trained models were applied to co-registered mpMRI volumes across the full tumor ROI. MRI volumes were skull-stripped using HD-BET, and whole-tumor segmentation was performed using a pretrained nnU-Net BraTS model, generating standard tumor compartment labels (enhancing tumor, non-enhancing/necrotic core, peritumoral edema) used for downstream spatial analyses. For spatial GCN inference, sliding-window patches were extracted across the tumor mask (excluding necrotic core for the NEU model), with patch-level feature values computed as mean signal intensity and standardized using training-cohort parameters. For each imaging feature, hierarchical clustering of patch values generated a multiscale event graph analogous to Section 3, and the trained two-layer GCN produced node-level subtype probabilities (softmax), which were aggregated to patch-level probabilities and projected back into image space via Gaussian kernel-weighted interpolation, followed by percentile normalization and smoothing.

Biological validity of MRI-derived subtype maps was assessed by comparing GCN-derived probabilities at biopsy locations with RNA-based subtype scores (logit-transformed), after removing tumor-specific effects within CE and NE regions separately; associations were assessed using Spearman correlation and linear mixed-effects models with patient-level random intercepts, including region-interaction terms. A combined NEU-minus-GPM subtype axis (logit scale) was similarly compared between MRI and RNA data. Patient tumors from the test cohort with complete imaging data (28/28 features) were included in these correlative analyses (8 patient tumors, 32 biopsies).

Finally, tumor-level NEU and GPM subtype burden was computed from voxelwise probability maps within manually defined NE and CE compartments, yielding compartment-specific and combined global subtype scores and a global metabolic dominance ratio. Overall survival was assessed in primary IDHwt HGG cases with available follow-up using Kaplan–Meier analysis (median split of the metabolic dominance score) and Cox proportional hazards models adjusting for age and NE fraction of tumor volume. Patients from the training and test cohort with complete imaging and clinical data were included in these outcome analyses (25 patients).

## Data Availability

All datasets analyzed in the current study will be made available upon reviewer and editor request and for publication along with this paper. Sequencing data will be deposited in a publicly accessible database. The remaining data are available within the Article, Supplementary Information and will be provided in the Source Data file.

## Code Availability

Code for all modeling analyses will be made available on Github upon reviewer or editor request and for publication along with this paper.

## Acknowledgements

We express gratitude for current and past members of the image analysis and glioma biopsy protocol teams. This work was supported by NIH grant U01CA220378 (N.L.T., L.S.H.); NIH grant U01CA250481 (L.S.H., N.L.T.); NIH grant R01CA264992 (L.S.H.); NIH grant R01CA221938 (L.S.H.); NIH grant R21NS082609 (L.S.H, N.L.T.); Center for Individualized Medicine Mayo Clinic (L.S.H., N.L.T.); The Medical Scientist Training Program at Mayo Clinic (5T32GM065841-20; M.J.F.)

## Author Contributions–

Conceptualization: MJF, SS, KK, NLT, LSH. Methodology: MJF, SS, GML, GM, JC, YZ. Software: MJF, GML, KK. Formal Analysis: MJF, SS, GML. Investigation: MJF, SS, MK, IH, BRB, AT, CK, RZ, JP, JL, KD, KB, LCB, CCQ, AI, SFE, MC. Resources: NLT, LSH. Data Curation: MK, IH. Writing: MJF. Review and Editing: KK, NLT, LSH. Visualization: MJF. Supervision: KK, NLT, LSH. Project Administration: AI, MC, KK, NLT, LSH. Funding Acquisition: NLT, LSH.

---

**Supplementary Figure 1.**
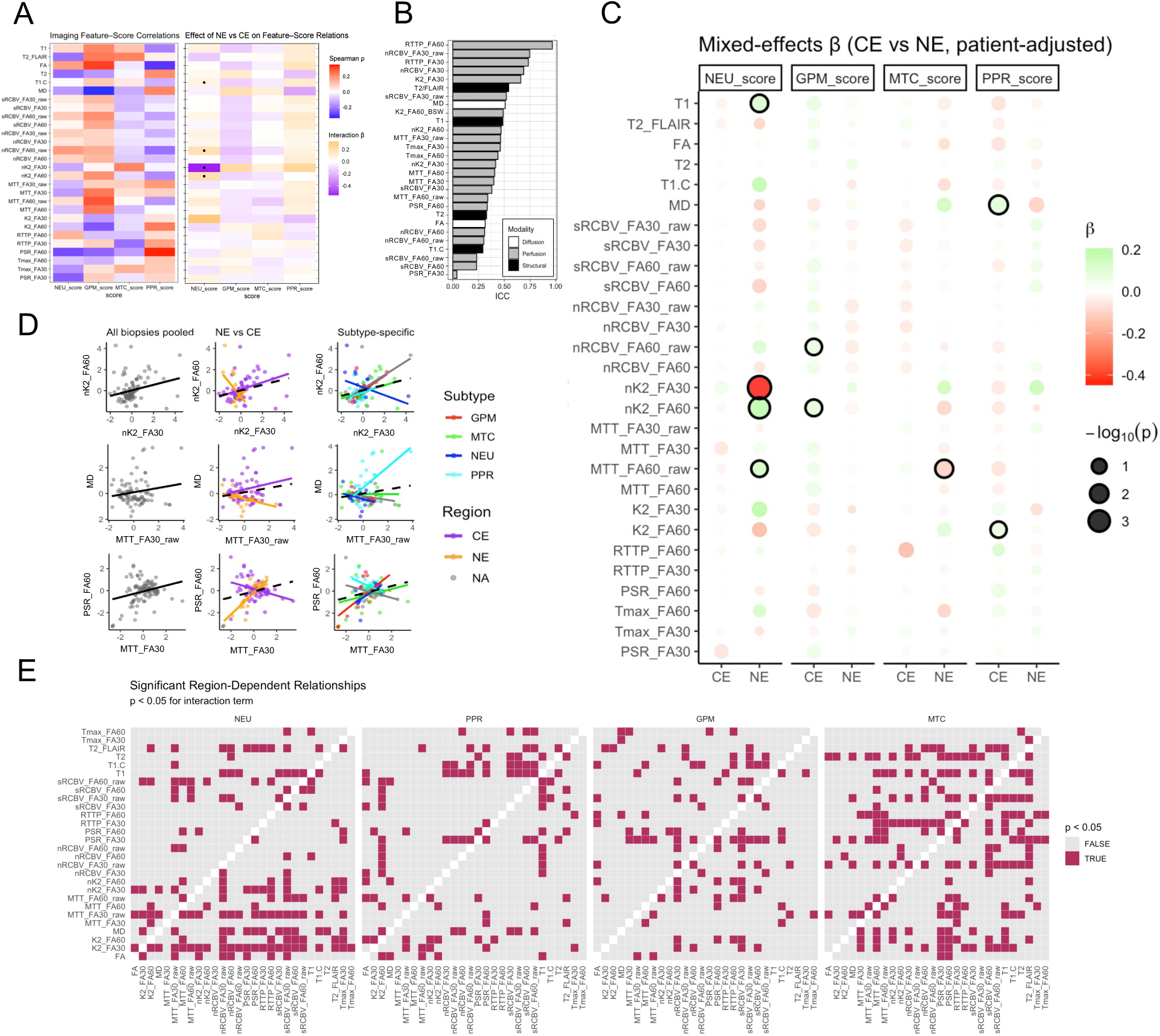
mpMRI signatures display region- and subtype-specific patterns. **A.** Spearman correlation of imaging features with pathways-based subtype scores (left). Interaction size reflecting directional shift in relation between imaging feature and subtype score in NE relative to CE samples (right). Significance (p<0.05) denoted with a dot. **B.** Patient-level intraclass correlation coefficient (ICC) per imaging feature quantifying extent of patient-effect per imaging feature. **C.** Linear mixed-effect model effect size for relations between imaging features and subtype score by tumor region (NE/CE) controlling for patient effects. Significant feature-subtype score relations (p<0.05) denoted with a black outline. **E.** Linear mixed effect models testing regional (NE/CE) interaction for paired imaging feature relations separated by molecular tumor subtype while controlling for patient effects. Significance (p<0.05) reflects strong influence of tumor region (NE/CE) on the absolute strength of linear relation between paired imaging feature signals at the biopsy level.

**Supplementary Figure 2.**
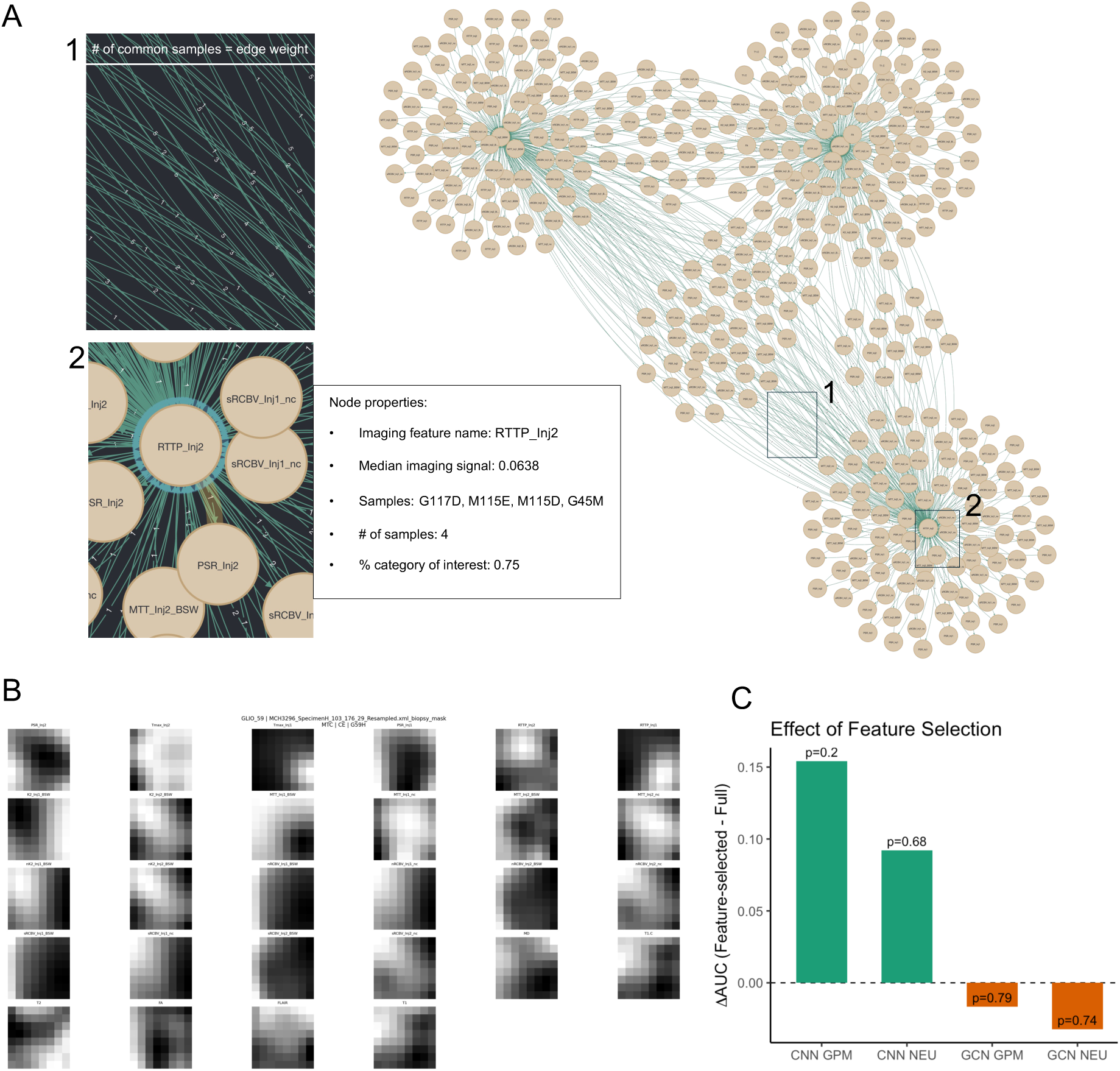
Graph network framework and convolutional neural network channels. **A.** Example segment of mpMRI-derived graph structure representing nodes generated through hierarchical clustering pipeline. Varied number of common samples between nodes represented edge weight (inset #1), leading to a dynamically-weighted multiresolution graph structure. Key node properties (inset #2) highlighted for a single node. “% category of interest” is equivalent to percentage of NE sample composition (for community detection) or percentage of NEU/GPM sample composition (for GCN). **B.** Example of 8x8 mpMRI channels for a single biopsy region of interest (biopsy G59H, MTC subtype, CE region) that served as an input for convolutional neural network prediction of molecular subtype. **C.** Change in AUC between GCN and CNN with and without prior MRMR feature selection using DeLong test. Feature name translation; Inj1 = FA60, Inj2 = FA30.

**Supplementary Figure 3.**
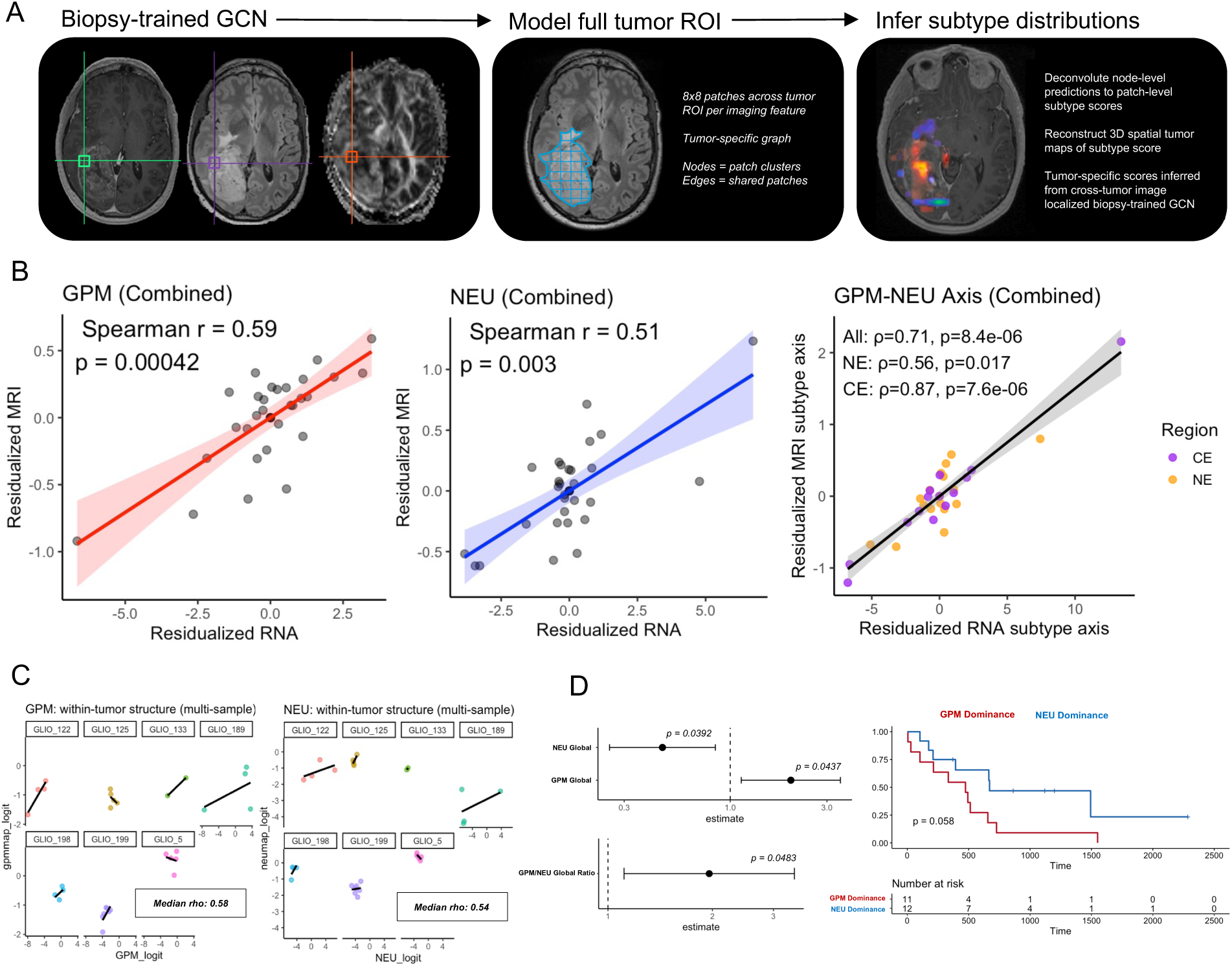
Tumor-specific subtype inference with biopsy-trained GCN. **A.** Pipeline for generation of tumor-specific GCN-inferred molecular subtype maps. **B.** Spearman correlations of molecular subtype map scores (right = GPM, middle = NEU, right = GPM-NEU subtype axis) with ground truth molecular subtype or subtype axis score across held out test tumors (n = 32 biopsies, 8 patient tumors). **C.** Within-tumor trend lines for raw subtype map score with ground truth molecular subtype score for held out test tumors comprising at least 2 biopsies. Median rho represents the median Pearson correlation across multi-sample tumors. **D.** Cox proportional hazards model assessment of associations between GCN-derived combined NEU enrichment (NEU Global), combined GPM enrichment (GPM Global), and the global GPM/NEU dominance ratio with overall survival, adjusting for age and non-enhancing tumor volume fraction (n = 23 GBM patients, left). Kaplan–Meier analysis stratified tumors by median global GPM/NEU dominance (n = 23 GBM patients, right).

