## Supplementary Methods for "Graph-based modeling of multiparametric MRI deciphers molecular states of high-grade glioma invasion with prognostic implications"

### **S1. Patient Cohort, Data Acquisition and Transcriptome Analysis**

This study was conducted under Institutional Review Board (IRB) approval from Barrow Neurological Institute (BNI) and Mayo Clinic Arizona, with the Mayo Clinic IRB serving as the overarching coordinating protocol. Written informed consent was obtained from all participants prior to enrollment. All data collection and protocol procedures followed approved guidelines and regulations outlined in the Mayo Clinic and BNI IRB protocols. Analyses were restricted to treatment-naive primary IDH-wildtype HGG (29 GBM, 6 other HGG. Samples with known technical or quality-control exclusions were removed prior to statistical modeling, yielding 134 image-localized biopsies from 35 primary tumors for the primary analysis cohort.

**S1.1 Imaging protocol, Multiparametric MRI acquisition**

Preoperative multiparametric MRI (mpMRI) was performed on 3-T clinical scanners within several days prior to surgery. Imaging protocols included conventional anatomical sequences (post-contrast T1-weighted (T1+C), T2-weighted (T2), and T2/FLAIR), diffusion tensor imaging (DTI), and dynamic susceptibility contrast (DSC) perfusion MRI. Quantitative imaging maps generated from these acquisitions included fractional anisotropy (FA), mean diffusivity (MD), relative cerebral blood volume (rCBV), and additional DSC-derived hemodynamic metrics. MRI acquisition parameters and image processing procedures followed the methodology previously described in detail in Hu et al. [4].

During image-guided surgery, neurosurgeons obtained multiple stereotactically localized biopsies from spatially distinct tumor regions, including both contrast-enhancing (CE) and non-enhancing (NE) tumor compartments. Biopsy coordinates were recorded intraoperatively using stereotactic navigation systems and subsequently coregistered to preoperative MRI datasets. Tissue specimens were collected from anatomically separated locations throughout the tumor to capture regional biological heterogeneity. Image preprocessing and spatial registration were performed according to the pipeline outlined in Hu et al. [4]. Briefly, non-quantitative MRI sequences underwent intensity normalization and bias-field correction, while all imaging datasets were coregistered to a common anatomical reference space. Quantitative parametric maps were generated using established processing workflows for DTI and DSC-MRI data [10, 28, 35, 37, 38, 56-58].

For each biopsy location, a spatially matched region of interest (ROI) measuring 8 × 8 × 1 voxels (approximately 9.6 × 9.6 × 3 mm) was centered on the recorded stereotactic coordinates. ROI placement was reviewed by an experienced neuroradiologist (L.S.H), and each biopsy was annotated as originating from either CE or NE tumor. Biopsies affected by substantial imaging artifact, necrosis, resection cavity, or other factors expected to compromise quantitative imaging measurements were excluded from analysis. Raw mean signal intensity values were extracted from each ROI across all imaging modalities and quantitative parametric maps. These biopsy-localized imaging measurements were subsequently used for correlative analyses and model development.

**S1.2: Sequence alignment and classification** RNA sequencing reads were aligned to GRCh38 using STAR (v2.7.0b), and gene-level counts were generated with featureCounts (v1.6.3). Raw counts were GC-content normalized using EDAseq (v2.22.0) and batch-adjusted using ComBat-seq. Transcriptional subtypes were assigned via the GBMclassifier Shiny application (<https://lucgar88.shinyapps.io/GBMclassifier/>), with each sample receiving the subtype with the highest significant enrichment score (logit NES > 0.40, p < 0.05).

### **S2. Imaging feature–biology association analysis**

**2.1 Regional interaction analysis (CE vs. NE).** For each imaging feature–subtype score pair, ordinary least squares regression models were fit according to Y=β0+β1+β2R+β3(X×R)+ε, where Y denotes the subtype enrichment score, X denotes the z-score-standardized imaging feature, and R denotes anatomical region (contrast-enhancing [CE] or non-enhancing [NE], with CE as the reference level). The interaction coefficient (β3) quantified region-dependent changes in the association between imaging features and subtype scores, with positive coefficients indicating stronger feature–score coupling in NE relative to CE regions. Interaction p-values were adjusted for multiple testing using the Benjamini–Hochberg false discovery rate (FDR) procedure and visualized as heatmaps.

**S2.2 Region-stratified correlation analysis** Spearman correlations between imaging features and subtype scores were computed separately within CE and NE samples (two-sided tests within each compartment). The difference in correlation strength was calculated as Δρ = ρ_NE − ρ_CE; feature associations showing directional reversal between compartments were classified as sign-flip events.

**S2.3 Mixed-effects modeling of imaging–molecular associations** Linear mixed-effects models were fit per feature–score pair as y_ij = β₀ + β₁X_ij + β₂R_ij + β₃(X_ij × R_ij) + b_i + ε_ij, where b_i is a patient-level random intercept and R_ij denotes region. CE effects correspond to β₁; NE effects correspond to β₁+β₃. Models were fit by REML, with fixed-effect significance assessed via Wald tests (lmerTest).

**S2.4 Patient-level variance (ICC) analysis** ICCs were computed per imaging feature using random-intercept mixed-effects models: ICC = σ²_patient / (σ²_patient + σ²_residual). Features were grouped into structural, diffusion, and perfusion modalities for comparative visualization.

**S2.5 Global region-dependent feature–feature interaction analysis** For each ordered imaging feature pair (excluding identical pairs), mixed-effects models were fit as Y_ij = β₀ + β₁X_ij + β₂R_ij + β₃(X_ij × R_ij) + b_i + ε_ij. β₃ quantified whether feature–feature coupling differed between CE and NE (positive = stronger NE coupling). Models used REML/lmerTest; p-values were FDR-corrected (Benjamini–Hochberg) and visualized as symmetric feature–feature heatmaps, with significant interactions (q<0.05) additionally shown as binary FDR-thresholded matrices.

**S2.6 Subtype-dependent feature–feature interaction analysis** For each ordered feature pair, mixed-effects models were fit as Y_ij = β₀ + β₁X_ij + β₂S_ij + β₃(X_ij × S_ij) + b_i + ε_ij, where S_ij denotes transcriptional subtype. ANOVA on the fitted model evaluated the global significance of the feature-by-subtype interaction (F-statistic and p-value). Models were fit using lme4/lmerTest (REML); p-values were FDR-corrected across all pairs and visualized as feature–feature heatmaps of −log₁₀(p), with significant interactions (q<0.05) annotated.

### **S3. Community Detection**

**S3.1 Imaging feature preprocessing** Near-zero-variance features were removed using frequency and uniqueness thresholds implemented in the caret framework. Highly collinear features were removed using pairwise Pearson correlation filtering at a threshold of 0.98. Remaining features were z-score normalized using cohort-wide feature means and standard deviations.

**S3.2 Hierarchical imaging event generation** For each imaging feature, Euclidean distance matrices were computed across samples and hierarchical agglomerative clustering performed using median linkage. Dendrogram partitions corresponding to all internal clustering events were extracted via recursive tree partitioning. For each event, the following were recorded: constituent sample identifiers, median imaging signal across included samples, imaging feature identity, and regional composition (percentage NE vs. non-NE samples). Both terminal sample-level clusters and higher-order merged states were retained.

**S3.3 Graph database construction** Events were imported into Neo4j using the py2neo interface, with each event represented as a node carrying attributes for median signal, constituent samples, sample count, regional composition, and event metadata. Two event nodes were connected if they shared at least one constituent sample, with edge weight equal to the number of shared samples.

**S3.4 Graph community detection and consensus analysis** Root nodes (containing all samples) and terminal leaf nodes (single samples) were excluded from community detection. For each of 1,000 iterations, one eligible node per imaging feature was randomly selected and assigned an iteration-specific label; a graph projection was generated using only nodes from that iteration, with relationships treated as undirected and weighted by the num_common_samples edge property. Community detection was performed using the Neo4j GDS Louvain modularity optimization algorithm:

(gds.modularityOptimization.stream, relationshipWeightProperty = "num_common_samples")

For each iteration, samples within a community were collapsed into a unique sample set, and contributing imaging modalities were recorded. Community assignments across all iterations were merged into a consensus community table.

**S3.5 Community-level regional enrichment and feature aggregation** For each consensus community, percentage NE and CE composition was calculated from biopsy-level regional annotations. Community-level imaging signatures were generated by aggregating z-score-normalized biopsy-level features using the median value across constituent samples. Communities with fewer than three samples or fewer than two unique imaging modalities were excluded.

**S3.6 Highly invasive community identification and clustering** Communities exceeding the 75th percentile of the global percentage-NE distribution were defined as highly invasive. Communities with identical constituent sample sets were collapsed to unique representatives prior to clustering. PCA was performed on standardized community imaging matrices, retaining components explaining ≥85% cumulative variance; Euclidean distance matrices in PCA space were clustered using Ward's minimum variance method (ward.D2). Cluster number was evaluated using average silhouette width across k = 2–10; a two-cluster solution was selected based on silhouette structure and biological interpretability, and visualized via PCA projection.

### **S4. Imaging Community–Transcriptomic Integration**

**S4.1 Redundancy reduction and pathway prioritization** Semantic similarity matrices among enriched GO-BP pathways were computed using the Rel similarity metric (rrvgo). Pathways were collapsed using a similarity threshold of 0.7, retaining representative parent terms with the strongest enrichment significance. Top pathways for each imaging cluster were prioritized based on adjusted p-value and normalized enrichment score (NES).

**S4.2 Differential imaging features between imaging-derived community clusters** Community-level imaging matrices were generated by aggregating standardized imaging features across all samples within each community using median values. For each imaging feature, between-cluster differences were assessed using two-sided Wilcoxon rank-sum tests; mean values within each cluster were calculated, along with directional (Cluster 1 − Cluster 2) and absolute differences. Multiple testing correction across features was performed using Benjamini–Hochberg FDR; features with FDR-adjusted p < 0.05 were considered significant. Cluster-specific imaging phenotypes were visualized using mean scaled feature plots and ridge density plots, with features ordered by absolute inter-cluster difference.

**S4.3 Association between imaging community clusters and NE enrichment** Differences in NE enrichment between clusters were evaluated using two-sided Wilcoxon rank-sum tests, with cluster-specific NE proportion distributions visualized using violin and box plots with overlaid individual communities, to assess whether distinct imaging-derived community states corresponded to differential enrichment of infiltrative tumor regions.

### **S5. Graph Convolutional Network Modeling**

**S5.1 Imaging feature preprocessing** Feature matrices were generated independently for training and test cohorts, split at the tumor level. Preprocessing followed the near-zero-variance filtering, Pearson correlation filtering (|r| > 0.98), and z-score normalization pipeline of S3.1. Missing values were imputed using median imputation calculated from the training cohort only and applied identically to the test cohort; z-score scaling parameters were similarly derived from training data only and applied to the test cohort. 28 of 28 imaging features were retained following filtering.

**S5.2 Feature selection (mRMR)** The mRMRe R package was used independently for each task (NEU vs. non-NEU; GPM vs. non-GPM) on the training dataset, binarized into one-vs-rest classification problems. Only numeric imaging-derived features were retained, missing values were removed, and the number of selected features was set to 28, ranked by mutual information criteria. For NEU, top-ranked features included FA, FLAIR, MTT_Inj2_nc, nK2_Inj1_BSW, nK2_Inj2_BSW, PSR_Inj2, RTTP_Inj1, RTTP_Inj2, T1.C, and Tmax_Inj2. For GPM, top-ranked features included FA, FLAIR, nK2_Inj2_BSW, K2_Inj2_BSW, MTT_Inj1_BSW, RTTP_Inj1, RTTP_Inj2, T1.C, Tmax_Inj2, and T1.

**S5.3 Event-based graph construction** Molecular subtype annotations were derived from integrated pathway-based subtype labels. For the NEU model, samples were labeled "NEU" vs. "Other"; for the GPM model, samples were labeled "GPM" vs. "Other." Hierarchical event generation and graph construction followed S3.2–S3.3, with each node additionally annotated by molecular subtype composition percentage. Edges were defined by shared sample membership, weighted by num_common_samples.

**S5.4 Graph preparation for deep learning** Graphs were imported from Neo4j via py2neo. Binary node labels were assigned using a ≥90% subtype-purity threshold (NEU or GPM composition, as applicable); remaining nodes were labeled negative. Imaging feature names were encoded via LabelEncoder to enable learned embeddings. NetworkX graphs were generated from edge tables and converted to PyTorch Geometric objects (from_networkx). Each node feature vector comprised the median imaging signal and a learned feature-identity embedding. Edge weights were computed as w_ij = log(1 + n_ij), where n_ij is the number of shared samples between connected nodes, then normalized by total unique samples and scaled to the maximum edge weight within the graph.

**S5.5 GCN architecture** The model included a learnable embedding layer for imaging feature identities, two GCNConv layers with ReLU activation, and dropout between convolutional layers. Node embeddings and imaging signal values were concatenated prior to graph convolution: x_i = [s_i ∥ e_i], where s_i is the median imaging signal and e_i is the learned feature embedding. The final layer produced binary classification logits per node.

**S5.6 Training strategy and hyperparameter optimization** Training graphs were partitioned into training (70%), validation (15%), and internal test (15%) node subsets via stratified sampling; these are node-level splits within training tumors only, distinct from the independent held-out test cohort (§S5.8). Class imbalance was addressed using weighted cross-entropy loss from inverse class frequencies. A sample-consistency regularization term was added: L = L_CE + λΣ_i w_i(p_i − p̄)², where p_i is the node-level predicted probability, p̄ is the sample-level mean probability, and w_i is the inverse sample-membership weight. Hyperparameters (hidden layer dimensionality, learning rate, dropout fraction, embedding dimensionality, sample-consistency weighting coefficient, and weight decay) were optimized using Optuna with five-fold stratified cross-validation on combined training/validation nodes. Each trial trained for up to 300 epochs using Adam with gradient clipping; model selection used mean validation F1 across folds with threshold-optimized predictions.

**S5.7 Final model training and evaluation** Final models were retrained using optimal hyperparameters on combined training and validation nodes, with early stopping based on loss convergence (patience = 50 epochs). Checkpoints stored network weights, architecture parameters, optimization settings, class weights, random seed, and classification thresholds. Node-level probabilities were aggregated to sample-level predictions via mean node logit: ẑ_sample = (1/N)Σlog(p_i/(1−p_i)), transformed via sigmoid: P = 1/(1+e^(−ẑ_sample)). Performance was evaluated using ROC-AUC, F1, and threshold-based accuracy at node and sample levels.

**S5.8 Independent test graph inference** Test graphs from held-out test tumors were generated using the identical preprocessing and graph construction pipeline, with imaging feature encodings aligned to the training graph via the previously fitted LabelEncoder. Test graphs were converted to PyTorch Geometric objects preserving node attributes, edge weights, feature identities, and sample mappings. Saved model checkpoints were applied directly without retraining; node-level probabilities were generated via forward inference and aggregated to sample-level via mean-logit/sigmoid transformation as in S5.7, with performance assessed via ROC-AUC at node and sample levels.

### **S6. Full Tumor ROI Analysis**

**S6.1 Brain extraction (HD-BET)** For each exam, a T1-weighted non-contrast image was automatically identified via filename pattern matching and used as input to HD-BET. The resulting brain mask was applied voxel-wise to all imaging modalities within the same exam, and masked images were saved in NIfTI format for downstream segmentation.

**S6.2 Automated tumor segmentation (nnU-Net)** Multiparametric MRI studies were formatted into nnU-Net-compatible input directories and processed using the pretrained BraTS 2020 task configuration (Task082_BraTS2020) with the 3D full-resolution model (3d_fullres) via nnUNet_predict, generating standard BraTS labels for enhancing tumor, necrotic core, and peritumoral edema/non-enhancing tumor.

**S6.3 Whole-tumor spatial GCN inference mapping** Tumor segmentations were registered to imaging space to define voxelwise tumor masks; for the NEU model, necrotic core voxels (BraTS label 1) were excluded, while all tumor voxels were retained for the GPM model. Sliding-window patches (8×8×1 voxels, 50% in-plane overlap) were extracted and retained if ≥25% of voxels fell within the valid tumor mask; patch-level feature values were the mean signal intensity across tumor voxels, standardized using training-cohort means/SDs. For each imaging feature, hierarchical agglomerative clustering (average linkage, Euclidean distance) was performed across patch values, with each dendrogram merge represented as a graph node (patch set, median intensity, feature identity); nodes sharing patches were connected with edge weights proportional to shared-patch counts. A two-layer GCN with learned feature-identity embeddings produced node-level probabilities via softmax; patch-level probabilities were computed by averaging across all nodes containing a given patch. Voxelwise maps were reconstructed via Gaussian kernel-weighted interpolation (σ=2.0) centered on each patch, with overlapping contributions normalized by summed kernel weights. Probability maps were normalized per tumor using 5th/95th percentile values and smoothed with a Gaussian filter (σ=2.0).

**S6.4 Biological validation against transcriptomic data** For spatially matched biopsies with transcriptional profiling, GCN-derived probabilities (from voxelwise maps) and RNA-derived subtype scores were both logit-transformed. For each region (CE, NE separately), linear models with patient/tumor identifier as a categorical factor were fit to MRI- and RNA-derived scores, and residuals were retained; residualized CE and NE datasets were combined, and Spearman correlations assessed MRI–RNA associations for NEU and GPM independently. A combined subtype axis was defined as logit(NEU) − logit(GPM) for both RNA and MRI-GCN scores, residualized as above, and compared via Spearman correlation. Linear mixed-effects models (patient/tumor random intercept) further assessed MRI–RNA concordance, with MRI-GCN scores as dependent variables and RNA-derived scores as fixed effects (separately for NEU, GPM, and the combined axis); interaction models including region and an RNA-by-region term tested whether MRI–RNA relationships differed between CE and NE. Fixed-effect significance was assessed using the Satterthwaite approximation.

**S6.5 Multimodal CNN-based biopsy classification (NEU and GPM reference models)** For each biopsy, a single axial slice was selected at the centroid (z-axis) of the biopsy ROI bounding box, and 2D patches were extracted across all modalities, resized to 8×8 pixels via bilinear interpolation and per-patch z-score normalized; modalities with missing data were zero-filled. All modalities were stacked as input channels. The CNN architecture comprised two convolutional layers (3×3 kernels) with batch normalization and ReLU, followed by a spatial attention module (1×1 convolution with sigmoid gating), global adaptive average pooling, and a fully connected layer producing a single logit. Training used BCEWithLogitsLoss with class weighting (negative:positive ratio), Adam optimizer (learning rate 1×10⁻⁴, 50 epochs), and a patient-level train/test split preventing leakage. Performance was evaluated using AUC, accuracy, confusion matrices, and classification reports, with stratified analyses for CE and NE biopsy regions.

**S6.6 AUC comparison** CNN and GCN ROC curves on matched held-out samples were compared using paired DeLong tests, accounting for correlation between predictions generated on the same biopsies.

**S6.7 MRI subtype burden and survival analysis** NE (regional label 1) and CE (regional label 3) compartments were extracted from registered tumor masks using manually defined segmentation labels (distinct from automated BraTS outputs, §S6.2), excluding necrosis. Mean subtype probabilities were calculated within each compartment for NEU and GPM maps. Global subtype scores were defined as the sum of z-scored NE and CE burdens (e.g., GPM_global = z(GPM_NE) + z(GPM_CE)). A global metabolic dominance score was calculated as: GPM_NEU_ratio_ALL = (GPM_NE + GPM_CE) / [(GPM_NE + GPM_CE) + (NEU_NE + NEU_CE)]. Survival analyses were restricted to 25 primary IDHwt HGG and the subset of 23 GBM cases with clinical follow-up. Kaplan–Meier analysis used a median split of the global metabolic dominance score. Cox proportional hazards models evaluated survival associations with (i) global NEU and GPM burden and (ii) the continuous metabolic dominance ratio, with multivariable models adjusting for age at diagnosis and NE fraction of tumor volume. Hazard ratios and 95% CIs were visualized as forest plots
